# ExTraMapper: Exon- and Transcript-level mappings for orthologous gene pairs

**DOI:** 10.1101/277723

**Authors:** Ferhat Ay, Abhijit Chakraborty, Ramana V. Davuluri

**Affiliations:** La Jolla Institute for Allergy and Immunology, La Jolla, CA, 92037, USA; UC San Diego, School of Medicine, La Jolla, 92093, CA, USA; Stony Brook University, Department of Biomedical Informatics, Stony Brook, NY, 11794, USA

## Abstract

Access to large-scale genomics and transcriptomics data from various tissues and cell lines allowed the discovery of wide-spread alternative splicing events and alternative promoter usage in mammalians. However, evolutionary studies primarily focus on gene-level orthology relationships, which hinders the importance of transcript-level diversity. Between human and mouse, gene-level orthology is currently present for nearly 16k protein-coding genes spanning a diverse repertoire of over 200k total transcript isoforms. Here we describe a novel method, ExTraMapper, which leverages sequence conservation between exons of a pair of organisms and identifies a fine-scale orthology mapping at the exon and then transcript level. ExTraMapper identifies more than 250k exon, as well as 30k transcript mappings between human and mouse using only sequence and gene annotation information. We demonstrate that ExTraMapper identifies a larger number of exon and transcript mappings compared to previous methods. Further, it identifies exon fusions, splits, and losses due to splice site mutations, and finds mappings between microexons that are previously missed. By reanalysis of RNA-seq data from 13 matched human and mouse tissues, we show that ExTraMapper improves the correlation of transcript-specific expression levels suggesting a more accurate mapping of human and mouse transcripts. ExTraMapper also reports better transcript-level mappings compared to Ensembl orthology for the human proto-oncogene BRAF and its mouse ortholog as well as several other example genes with important isoform-specific functions. ExTraMapper is applicable to any pair of organisms that have orthologous gene pairs and is available at https://github.com/ay-lab/ExTraMapper and http://ay-lab-tools.lji.org/extramapper

## INTRODUCTION

The fundamental process of gene expression in mammalian genomes is a highly complex procedure that is regulated at multiple different levels (1). An important part of this regulation involves the transcription of alternative gene products from a single gene through the use of alternative promoters and alternative splicing (2-5). Alternative transcription allows an organism to produce multiple transcripts from a single gene, and in turn, these transcripts are translated into multiple protein isoforms that may substantially differ in their structures and functions (6-8). Recent advances in sequencing technology and the analysis of sequencing data now allow researchers to explore “one gene → multiple isoforms → multiple functions” paradigm instead of still commonly used “one gene → one protein → one function” approach (9-11). However, the common mode of research remains to be gene oriented partly due to informatics challenges posed by multiple and partially overlapping transcripts, including the very fundamental problem of quantification of isoform level expression (12).

While many bioinformaticians are making big strides in overcoming the challenges of transcript/isoform level gene expression analysis (12-15), only a few studies so far attempt to comparatively analyze transcripts to find transcript-level orthology relationship among different organisms (16,17). One important intermediate step in revealing transcript-level relationships between a pair of organisms is to identify their conserved exons. Again, in the literature, there is only a limited number of studies that aim to find exon-level relationships among different organisms (18-22). These methods use either sequence alignment tools, such as BlastN and tBlastX, or reverse engineer the protein level similarity between isoforms to find conserved exons between two organisms. Often, only the fully coding exons are considered by these methods completely leaving out the partially coding and the non-coding exons such as first and last exons of a transcript. At the transcript level, Exalign, described in Pavesi et al.(16) and used for transcript mappings in Zambelli et al. (17), computes similarities between two given transcripts by utilizing only the exon-intron structure and the coding lengths of exons. Exalign first calculates a pairwise exon similarity score that solely depends on the length difference between the two exons and then uses these scores to find the best local alignment between the exon-intron structures of the two transcripts. As we will discuss further, Exalign does not utilize any sequence similarity or orthology information between the exons or the transcripts and is biased towards assigning higher similarity scores for transcripts with large number of exons.

Here we describe a new method, *ExTraMapper*, which extracts fine-scale mappings between the exons and transcripts of a given pair of ortholog genes between two organisms using sequence conservation between the two genomes and their gene annotations **(Fig. 1)**. To demonstrate the use of our method, we find exon and transcript mappings for nearly 16k protein coding genes with orthology defined between human and mouse. Compared to previous methods, ExTraMapper identifies a significantly larger number of exon, as well as transcript mappings with differing stringencies for conservation. Notably, our exon mappings include microexons (< 21 bp), which are largely missed by previous approaches. ExTraMapper also provides a more detailed exon classification in terms of exon conservation status and exon loss/gain from one organism to the other. Using similarity both in coding domain and in overall sequence, ExTraMapper better distinguishes transcript pair similarities leading to resolution of many-to-many relationships between transcripts into one-to-one when possible. Furthermore, transcript similarity scores computed by ExTraMapper show less dependency to the number of exons of a transcript compared to an existing method that uses only exon length information for scoring transcript pairs. We also compared and found that RNA-seq expression profiles (23) of orthologous transcripts from human and mouse tissues identified by ExTraMapper show a higher degree of correlation and more similar extent of expression level then Exalign specific isoforms. Our results on multiple important human-mouse ortholog gene pairs and comparing their respective isoform-level expression profiles across different tissues from human and mouse show that ExTraMapper finds biologically relevant transcript-level mappings. The results also provides examples of exon fusions, splits, and losses due to splice site mutations.

**Figure 1.**
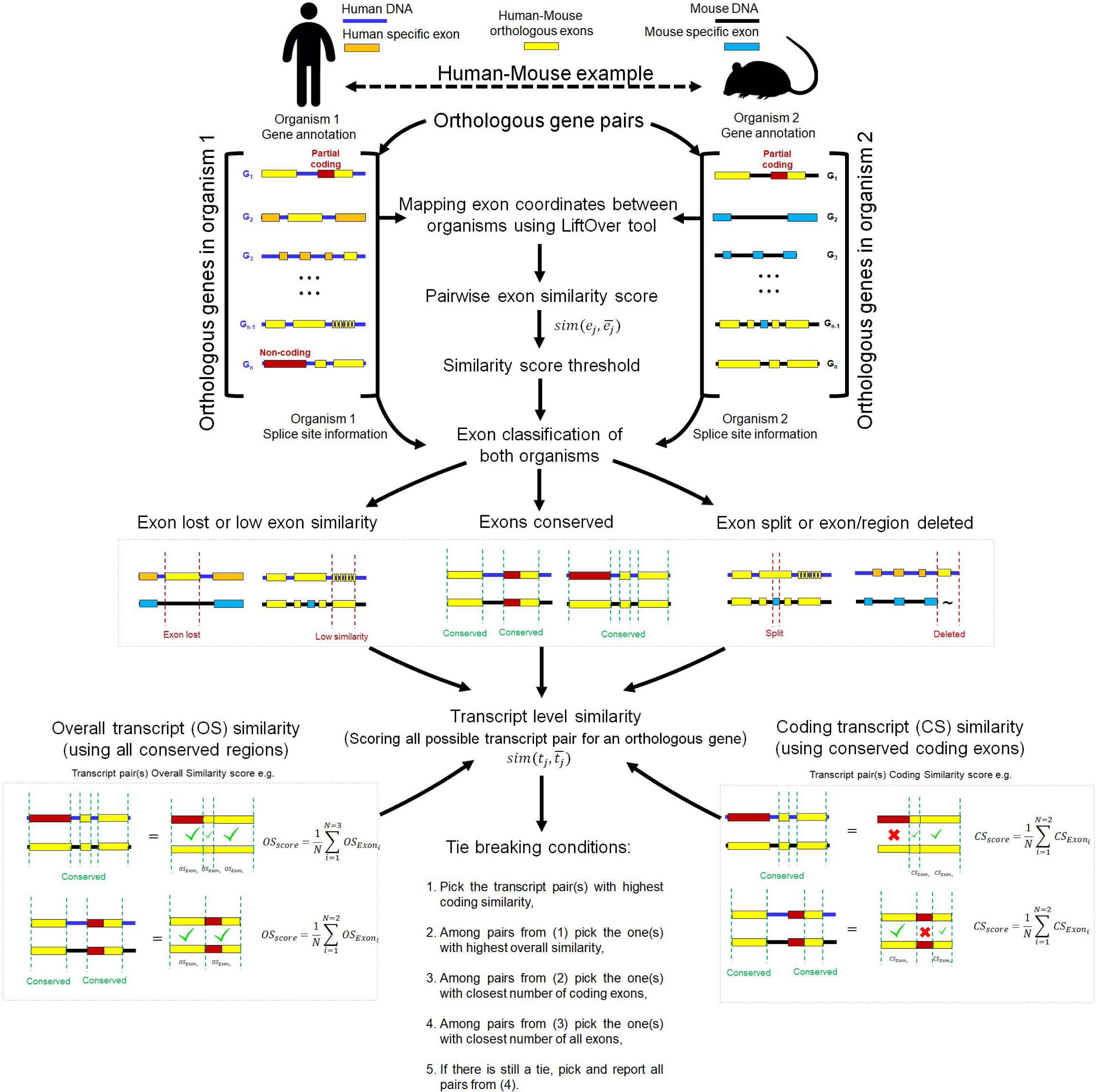
Overview of ExTraMapper. For each orthologous gene pair between the two organisms, ExTraMapper first computes exon-level similarities using gene annotations and the liftover tool that maps regions from one organism to the other. Each exon is then categorized according to its conservation status using the computed similarity scores and splice site information from acceptor and donor sites. The exon-level similarity scores are used to compute transcript-level similarities through an exon pairing algorithm. The transcript-level mappings for each gene are then determined using a greedy method that applies multiple tie break conditions to ensure mostly one-to-one mappings.

## RESULTS

### ExTraMapper utilizes exons with different coding and positional types

An important step in expanding orthology from gene level to transcript isoform level is to establish an orthology relationship between the exons, which are the building blocks of transcripts. To define such a relationship, we gather all exons from 15,846 gene pairs that are in human-mouse orthology and that have both genes annotated as protein coding (24) **(Methods)**. This provides us with nearly 500k and 345k exons for human and mouse, respectively. We note that Ensembl annotates the same exact coordinates with multiple different exon IDs when the same coordinates show varying coding activity across different transcripts. Collapsing such duplicate coordinates in one single entry, we get around 407k and 289k unique exon coordinates, respectively. **Figs. 2a, d** plot the distribution of the number of all exons and coding exons per gene for human and mouse genomes, respectively. The median numbers of exons and coding exons per gene were 22 and 14 for human, and 15 and 11 for mouse.

**Figure 2.**
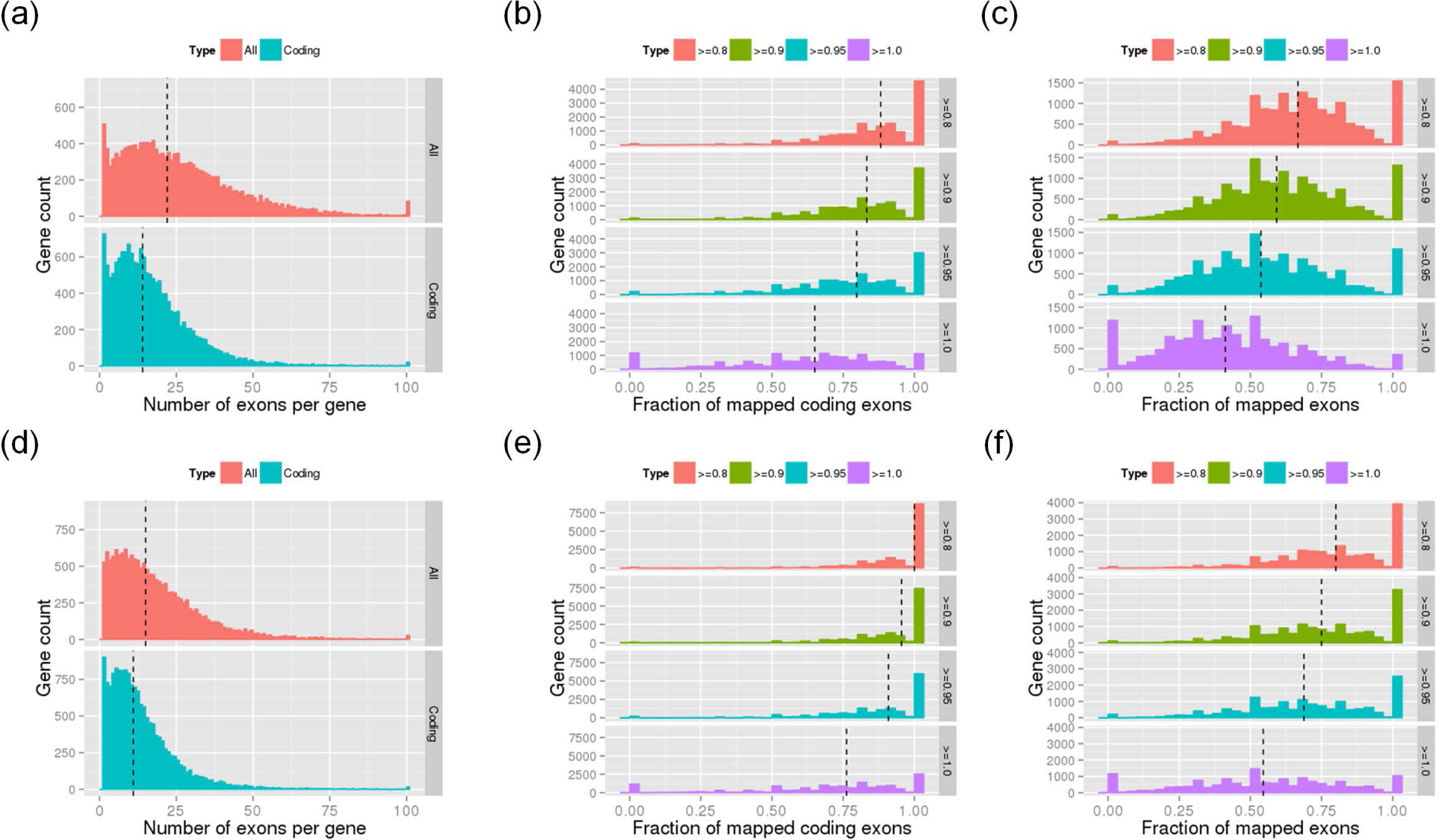
Summary of exon-level statistics and ExTraMapper mapping results for human (top row) and mouse (bottom row). **(a**,**d)** The histograms of the number of all and coding exons per gene. **(b**,**e)** The histograms of the fraction of coding exons (fully coding or part-coding) that are mapped when different stringency cutoffs for mapping similarity are used. A fraction of 1 indicates all exons of a gene are mapped and 0 indicates none are mapped. **(c**,**f)** Similar histograms using all exons (fully coding, part-coding, non-coding). Dashed vertical lines correspond to median of the distribution for each plot.

**Supp. Table 1** reports further grouping of these unique coordinates with respect to their coding types (non-coding, fully coding or partially coding) and their positions within each transcript they are involved (first, middle, last). Note that for each classification, namely coding type and position; we report a fourth category for exons that have multiple different types in different transcripts. Our further analysis of different exon types reported in **Supp. Table 1** using evolutionary conservation scores demonstrate that coding exons are more conserved compared to non-coding exons as expected **(Supp. Fig. 1)**. Also, exons that are exclusively in the middle of each transcript they participate are highly conserved **(Supp. Fig. 2)** and are mainly fully coding (43.7% and 60.5% for human and mouse) fully coding. In comparison, full coding exons make up only 10.7% and 8.7% among first or last exons of human and mouse genomes, respectively. These numbers are in agreement with the commonly used transcript or gene structure models with part-coding or non-coding first and last exons and fully coding exons in between them. However, these numbers also suggest that a considerable fraction of the human and mouse transcripts may not follow such a transcript model, which is assumed by several existing methods for exon or transcript level matching (16,17).

### ExTraMapper identifies conserved exons between human and mouse genomes

Without any assumption on the gene or transcript structure, ExTraMapper computes exon-level similarity scores and classifies exons with respect to their conservation between human and mouse (Methods). **Fig. 2** summarizes the fraction of exons for each gene that ExTraMapper mapped from human to mouse **(a-c)** and from mouse to human **(d-f)** with different stringencies of conservation as measured by the overlap after lifting over the coordinates. We compute this fraction for coding exons (fully or part coding) and all exons (including non-coding) separately. With an exon similarity threshold of 0.9, more than half the genes had at least 83% (human) and 96% (mouse) of all their coding exons mapped **(Figs. 2b, e)**. These percentages are 59% (human) and 75% (mouse) when all exons are considered **(Figs. 2c, f)**. Notably, a larger fraction of mouse exons is classified as mapped for each threshold choice reflecting the smaller number of total exons for mouse compared to human.

We also compute the inclusion level for an exon as the fraction of all protein coding transcripts that include a given exon (21, 25), and ask whether the inclusion level is a determinant of exon conservation. Similar to Modrek et al. (21), we observe that very large percent (95.9% for human and 95.7% for mouse) of constitutive and major-form exons (i.e., inclusion level >0.5) are mapped between the two organisms by ExTraMapper. For minor-form exons (i.e., inclusion level <0.5) the percentages of mapped ones are 52.6% and 58.8% for human and mouse, respectively. These results are in conjunction with Modrek et al. which suggests that around 98% of constitutive and major-form exons and only around 28% of minor-form exons existed before the divergence of the mouse and human genomes (21). The difference in the percentages between Modrek et al. in 2003 (∼40k total exons) and our work here (>840k total exons) is likely due to improvement in the exon annotations of both organisms within the past decade.

### ExTraMapper reports additional exon mappings compared to existing methods

We compare ExTraMapper with two previous methods that find exon-level mappings between human and mouse. For an orthologous gene pair, OrthoExon uses a two-step Blast search to find orthology exon pairs (20). OrthoExon first performs BlastN alignment from all human exons to mouse, and vice versa, to find significant hits (20). In the second step, OrthoExon realigns the exons with no significant hits from one organism to similar exons of the other using tBlastX to allow for sequence divergence. There are several issues with OrthoExon due to use of alignment-based scores for finding exon mappings. First, sequence alignment scores are more significant for longer exons leading to length bias when thresholded to determine “alignment hits”. Second, a Blast hit does not necessarily imply orthology and the hit score is not interpretable in terms of the percent overlap between the two aligned exons. Another exon mapping method by Zhang et al. uses protein level information from Inparanoid database to find ortholog and in-paralog exons between human and mouse using (22). This approach excludes first and last exons and requires exact length match between the two exons to deem that they are orthologs. These criteria substantially limit the number of exons considered for mapping and do not allow any sequence divergence further limiting the number of relationships that can be found between human and mouse exons. Furthermore, neither of the two methods above provide any information about how the exons that do not have orthologs diverged between the two organisms.

For comparison with our exon mapping results, we download and process the exon mappings from the two methods mentioned above to have all exon IDs in Ensembl format and all exon coordinates in hg38 and mm10 reference genomes (Method). **Fig. 3a** provides a summary of the resulting comparison of these methods with ExTraMapper run using multiple similarity thresholds for exon classification. ExTraMapper reports a significantly larger number of exons that are ***conserved*** for both organisms even for the most stringent threshold of 1.0, which means 100% overlap between the two exons when one is lifted over to its coordinates in the other organism. One main reason for this improvement is that ExTraMapper does not discard part-coding exons or first and last exons and uses both coding similarity and overall similarity to find mappings. Indeed, ExTraMapper finds perfect mappings (threshold of 1.0) for more than three times as many part-coding exons compared to two previous methods. Among the 78,037 part-coding human exons, for 53,231 ExTraMapper reports perfect mappings more than 38k of which were not reported by other methods. Similar numbers hold for mouse genome with 36,109 ExTraMapper perfect mappings out of 50,933 part-coding exons. Another important advantage of ExTraMapper is that, since it uses sequence conservation information at the exact coordinate level, it allows identification of mappings between very short exons, termed microexons (26), between human and mouse. Microexons are known to be highly conserved and have recently been shown to encode for protein domains involved in protein-protein interactions in the context of neurogenesis and autism (27-29). To assess the ability of different tools in capturing orthologous mappings of microexons, we compared ExTraMapper, OrthoExon, and Inparanoid. When we define microexons as exons that are shorter than 21 bps, ExTraMapper mapped 939 human and 829 mouse microexons, whereas OrthoExon and Inparanoid reported <100 and <350 exons, respectively **(Fig. 3B)**. Similar trends hold when we use a more liberal threshold of <51 bps for microexon definition with ExTraMapper reporting ∼4-6k additional exon mappings while capturing nearly all mappings reported by the other two tools **(Supp. Fig. 3)**.

**Figure 3.**
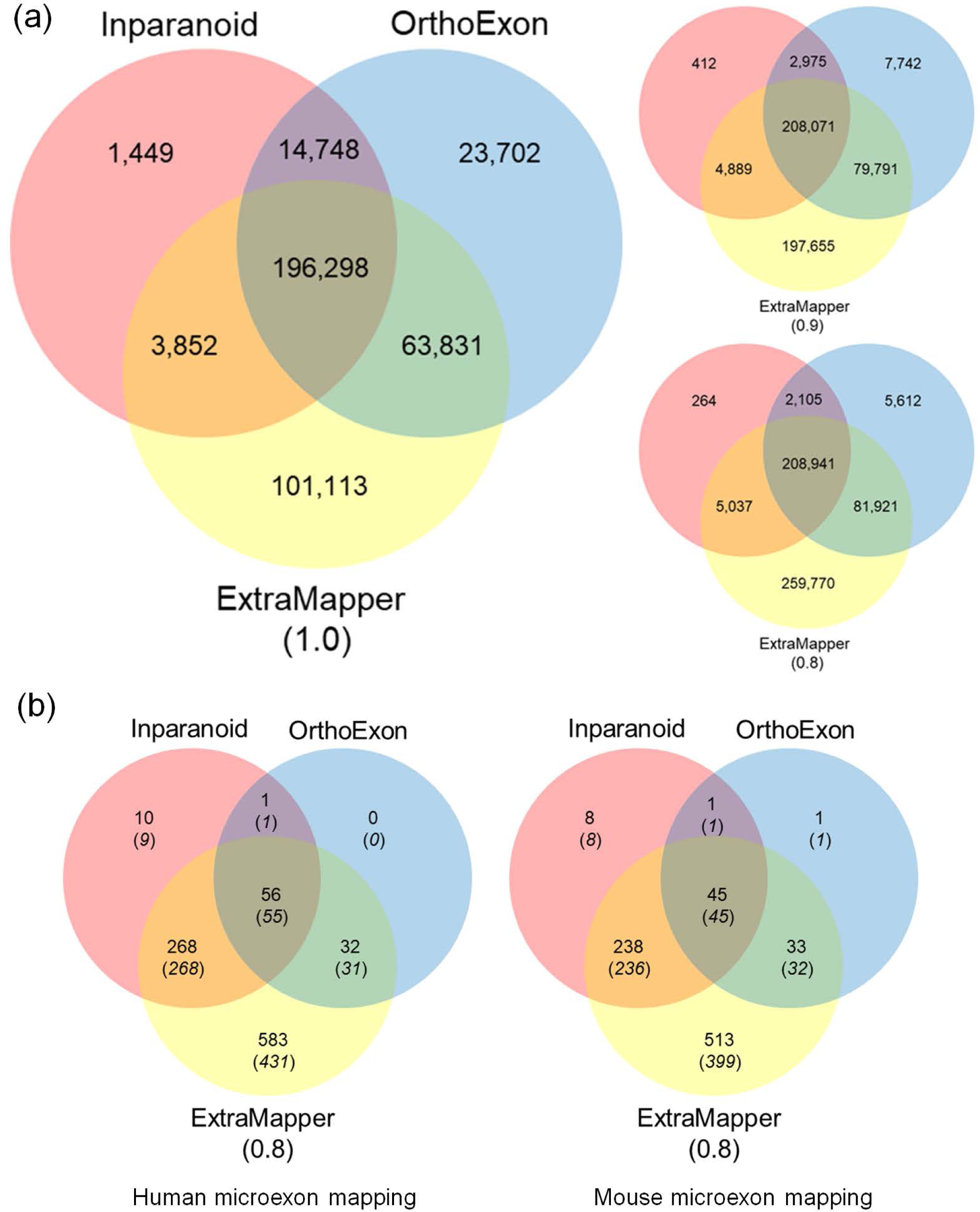
Summary of exon-level mapping results from three different computational approaches for human and mouse genome. Venn diagrams showing intersections among the three different methods: ExTraMapper, OrthoExon and Inparanoid. **(a)** When all exons are considered. Results for different ExTraMapper exon similarity score thresholds (1.0, 0.9 or 0.8) are shown separately. **(b)** When only microexons (<21bp) are considered with an ExTraMapper exon similarity score threshold of 0.8. Results for the number of human (left) and mouse (right) microexons are shown separately.

ExTraMapper further classifies the remaining exons, which are not conserved into three subclasses. The first group, ***region lost***, includes exons for which there is no corresponding region once the coordinates are lifted over to the other organism. The exons in this group are completely deleted, partially deleted or split in the other organism. The set of exons that are completely deleted is independent of *liftOver* parameter choices (such as minMatch), whereas the other two categories can vary with respect to these parameter choices. The second group, ***region conserved but exon lost***, consists of exons whose coordinates are successfully lifted over but to regions that are not annotated as exons in the other organism. This group can be further divided into two with respect to conservation of acceptor (AG di-nucleotide) and donor (GT di-nucleotide) sites in the lifted over coordinates. Some exons lose one or both of these splice sites rendering loss of exon function, whereas others preserve these sites but still are not annotated as exons in the other organism. The third group are the remaining non-conserved exons which were successfully lifted over to some exon regions in the other organism but with an overlap below the desired similarity threshold to be deemed conserved or mapped. We do not further analyze exons that belong to this last group because the conservation information for such exons is ambiguous. Later in the text we will provide examples of exons that belong to all other groups (conserved, region lost, region conserved but exon lost) in our case study of the *BRAF* gene. Our further analysis also shows the significant differences between evolutionary conservation score distributions of different classes of exons as determined by ExTraMapper **(Supp. Fig. 4)**. As expected, mapped exons mostly have high conservation scores and the ones in region lost group have lower scores which more uniformly span the whole range and has a median around 0.5. Interestingly, the small group of exons with the region conserved but exon function lost show a bimodal distribution where some exons have very high and others have very low evolutionary conservation scores. Further analysis shows that such exons with low conservation scores correspond to those with changes in their acceptor or donor sites between the two organisms.

### ExTraMapper computes transcript-level similarities using exon-level conservation

Now that we compute exon similarity scores and classify human and mouse exons according to their conservation between the two organisms, the next step is to compute similarity scores and establish mappings at the transcript level. Ensembl annotations for the 15,846 gene pairs we analyze here provide nearly 126k and 69k transcripts out of which more than 68k and 40k are protein coding for human and mouse, respectively. For these genes, the human orthologs have a median number of 6 transcripts and 3 coding transcripts per gene, whereas these numbers were 3 and 2, respectively, for the mouse genome **(Supp. Fig. 5)**. The smaller number of transcripts for mouse in comparison with human is in agreement with the smaller number of exons per gene for mouse. Whether these numbers reflect differences in biological diversity of transcripts or differences between the annotation qualities of the two organisms, it is clear that the number of mouse transcripts will be the bottleneck for transcript mappings that will be identified by ExTraMapper or any other algorithm.

We compute transcript-level similarity scores using a greedy method to find non-overlapping and non-contradicting set of exon pairings for a transcript with that maximizes the sum of exon similarity scores, either overall or only coding **(Methods)**. Our motivation behind using a greedy approach instead of a sequence alignment-like dynamic programming approach is that we want to favor exact or near exact mappings of exons over multiple mappings of lesser quality. For instance, given the choice between two exon mappings one with a score of 0.6 and one with a score of 0.7 and a single perfect mapping with a similarity score of 1, we prefer the latter, which is a conserved exon between the two organisms. A greedy approach will pick this conserved pair whereas a dynamic programming approach will do vice versa. Using this greedy approach, we compute an overall similarity score and a coding similarity score for a total of 730,440 pairs of transcripts generated from the 15,846 orthologous gene pairs. There are only 13,701 and 70,737 transcript pairs (1.9% and 9.7% of all possible pairs) giving a similarity score of 1 and 0.9 respectively either from the whole transcript bodies or by their protein coding portions. These pairs have the same number of exons and have each of their exons conserved 100% and 90% respectively (exon similarity threshold of 1 and 0.9). Such transcript pairs span in total 36,464 and 29,389 different human and mouse transcripts out of which 16,585 (45.5%) and 16,937 (57.6%) appeared in more than one perfect pair, respectively, suggesting one-to-many mappings exist even within perfectly conserved sets of transcripts of the two organisms. This is partly due to multiple transcripts in Ensembl with the same exact set of coordinates and partly because a perfect conservation in coding similarity still allows for difference in non-coding regions such as untranslated regions (UTRs).

### ExTraMapper identifies mainly one-to-one transcript mappings between human and mouse

To reduce the number of one-to-many transcript mappings, we employ a set of tie breakers as outlined in **Fig. 1** and described in detail in **Methods**. Briefly, for a given gene pair with transcript similarities computed between all possible human-mouse pairs, we first identify the transcript pairs with the maximum coding similarity. If there is only one such pair, we report this pair as a mapping and remove each transcript from their corresponding transcript sets and repeat the search for the pair(s) with maximum coding similarity. In case of a tie, we then look at, in a particular order, the overall similarity, the difference in the number of coding exons, and the difference in the number of all exons among all pairs with maximum coding similarity. If none of these tie breakers work then, we report all such pairs allowing some number of one-to-many mappings that are equivalently good. Similar to the computation of transcript similarity scores, we choose to use a greedy approach because we want to favor a smaller number of highly conserved transcript pairs instead of a larger number of moderately conserved ones. After this greedy approach with tie breaks, we get 8,007, 25,146 and 30,388 transcript pairs with either coding or overall transcript similarity score of 1, greater than 0.9 and greater than 0.8, respectively. These transcripts span a total of over 15k human-mouse gene pairs. Out of these, only 177 human and 195 mouse transcripts are reported as one-to-many mappings suggesting that ExTraMapper eliminates a very large portion of one-to-many mappings when there is enough information to distinguishing between multiple pairs with equal coding similarity.

We next compare the transcript mappings from ExTraMapper with those from a previous method, namely Exalign, which uses fully coding exon lengths to compute transcript similarities (16,17). The total number of human transcripts that are exactly mapped to only one mouse transcript is 30,013 for ExTraMapper (score >0.8) while this number was only 10,405 for Exalign **(Supp. Fig. 6a)**. Accordingly, ExTraMapper only reports 177 human transcripts that map to more than one mouse isoform, whereas Exalign reports 13,462 such human transcript. A similar trend can also be seen when numbers are computed with respect to mouse transcripts **(Supp. Fig. 6b)**. The fraction of one-to-one mappings is obviously dependent on the score threshold for ExTraMapper, however, even for a more stringent threshold of 0.9, ExTraMapper still reports over 25k such human transcripts suggesting its utility for breaking ties among potential one-to-many or many-to-many mappings. To study this on a specific example, we analyze the retinoic acid receptor alpha (*RARa*) gene, which is a critical nuclear receptor expressing specific isoforms that are found either in nucleus or cytoplasm and perform distinct functions depending on their cellular compartment (30, 31). Even though six possible pairs among two human and three mouse isoforms for this gene result in perfect coding similarity scores (both from ExTraMapper and Exalign), ExTraMapper breaks ties using the similarity between non-coding portions of these transcripts to report two one-to-one transcript mappings **(Supp. Fig. 7)**. We also compare ExTraMapper and Exalign transcript mappings for three other important genes (*TP53, TP63* and *TP73;* **Supp. Table 2**), which are known to play multiple critical roles in cell differentiation and response to stress through their rich repertoires of isoforms (8). For instance, for the 24 protein-coding human transcripts of *TP53* and 4 protein-coding mouse transcripts of *Trp53*, Exalign reports 16 pairs with identical scores (among 4 human and 4 mouse transcripts). In contrast, ExTraMapper, using information from other exons including those that are part-coding, reports four different one-to-one mappings **(Supp. Table 2)**. Such differences between the two methods in distinguishing transcript pairs from each other is observed for other genes where multiple different transcripts have the same number of exons such (e.g., 11 exons for the case of *TP53*). For *TP63-Trp63 and TP73-Trp73*, since most isoforms vary in exon length both methods successfully capture transcript mappings **(Supp. Table 2)**.

### ExTraMapper eliminates biases in transcript similarity score computation

We compare the transcript similarity scores computed by ExTraMapper with those from Exalign, which uses fully coding exon lengths to compute transcript similarities (16, 17). For each possible length difference *d* between two exons, one from human and one from mouse, Exalign calculates the probability of finding two exons with length difference at most *d* and uses the minus log of this probability as their exon similarity score. These similarities are then used together with a gap penalty to perform a structural alignment between a pair of transcripts each of which is represented as a sequence of exon lengths. Each alignment reports (i) a Blast-like alignment score that is independent of the query or database size, and (ii) an expectation value (E-value) that corresponds to the number of equally good or better hits expected by chance when searching the query transcript against a database of a particular size. For our comparison with ExTraMapper, we use Exalign’s alignment scores which are perfectly correlated the minus log of E-values.

We download and apply Exalign to the same set of gene annotations used for ExTraMapper for a fair comparison (Methods). First, we compare the best Exalign alignment score and the best ExTraMapper similarity score for the top scoring transcript pair for each gene with Exalign E-value < 0.001 and ExTraMapper score ≥ 0.8 (either coding or overall transcript similarity). **Fig. 4a** shows that there is very strong correlation between the best Exalign score and the number of exons a human transcript has (Pearson’s r=0.94). On the other hand, ExTraMapper scores have substantially lower dependence on the transcript length (**Fig. 4b**, Pearson’s r=0.07). Both figures are in agreement that human transcripts which code longer proteins (over 50 coding exons) have very similar ortholog transcripts in mouse, suggesting they are evolutionarily conserved. However, a near-linear relationship between the transcript length and Exalign scores across the whole range of lengths suggest that the Blast-like score used by Exalign is highly biased by the transcript length. Similar plots with respect to mouse transcript lengths exhibit the same trend **(Fig. 4c,d**). We repeat the same analysis using Exalign and ExTraMapper transcript similarity scores for all transcript pairs (**Supp. Fig. 8;** rather than the best scoring one for each gene as in **Fig 4**) again, highlighting a stronger dependency of Exalign scores to transcript length compared to ExTraMapper.

**Figure 4.**
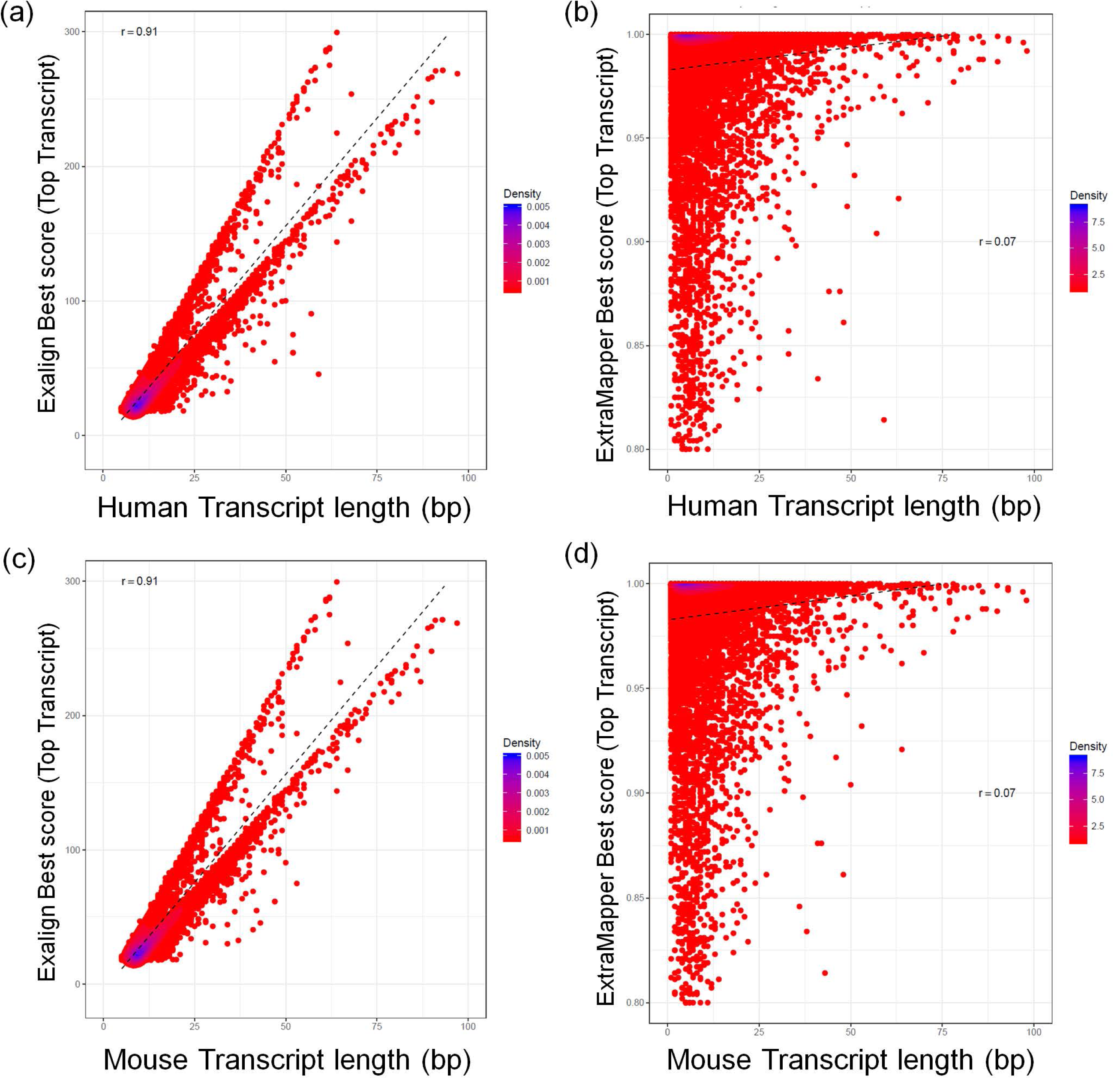
The length bias of the transcript-level similarity scores computed by Exalign. The density plots of the best Exalign alignment score and the best ExTraMapper similarity score for the top scoring transcript pair for each gene with Exalign E-value < 0.001 and ExTraMapper score > 0.8 (either coding or overall transcript similarity) are plotted for **(a)** human transcripts using Exalign **(b)** human transcripts using ExTraMapper **(c)** mouse transcripts using Exalign **(d)** mouse transcripts using ExTraMapper. Transcript length indicates the number of coding (either fully or partly) exons of a transcript. Pearson’s correlation is reported for each figure together with a linear fit.

### Expression profile comparison of orthologous transcript pairs

Mouse models are the bed stone of mechanistic studies of human genes and their role in different diseases due to conserved sequence, function and expression profiles of their orthologous genes (23,32-35). Previous work has shown that expression profiles of the conserved genes between the two organisms is highly similar between matched tissues (23,36) by reanalysis of human and mouse tissue-specific RNA-seq expression profiles (35). Here we use the same dataset **(Supp. Table 3)** to compare transcript-level expression estimates from human and mouse for pairs of transcripts identified by either ExTraMapper or Exalign. For reference, for each of the 13 tissues, we first compute the correlation between expression values at the gene level as was done by Gilad et al (36) resulting in a median R of 0.83 (Pearson correlation; **Fig. 5a**). We then repeat the correlation calculation using top orthologous transcript pairs obtained either from ExTraMapper, from Exalign or by both (common pairs). We observe that common transcript pairs exhibit the highest correlation followed by pairs predicted by ExTraMapper **(Fig. 5a)**. Exalign reported transcript pairs slightly lower correlation as well as higher quantile rank difference of individual transcripts based on their expression distribution in every human and mouse tissue compared to common pairs and pairs from ExTraMapper **(Fig. 5b)**. To better understand the source of higher expression correlation and smaller rank difference for ExTraMapper pairs compared to Exalign, we next compare the same entities using a subset of 562 human and 469 mouse transcripts where ExTraMapper and Exalign both identified a single orthologous isoform in mouse and human respectively, but the identified orthologous partners are different by both programs. For these 562 human and 460 mouse transcripts with their respective ExTraMapper and Exalign identified orthologous partners in the other organism, the transcript-level expression correlations also show marginally higher values for ExTraMapper mappings compared to Exalign **(Fig. 5c,d)**.

**Figure 5.**
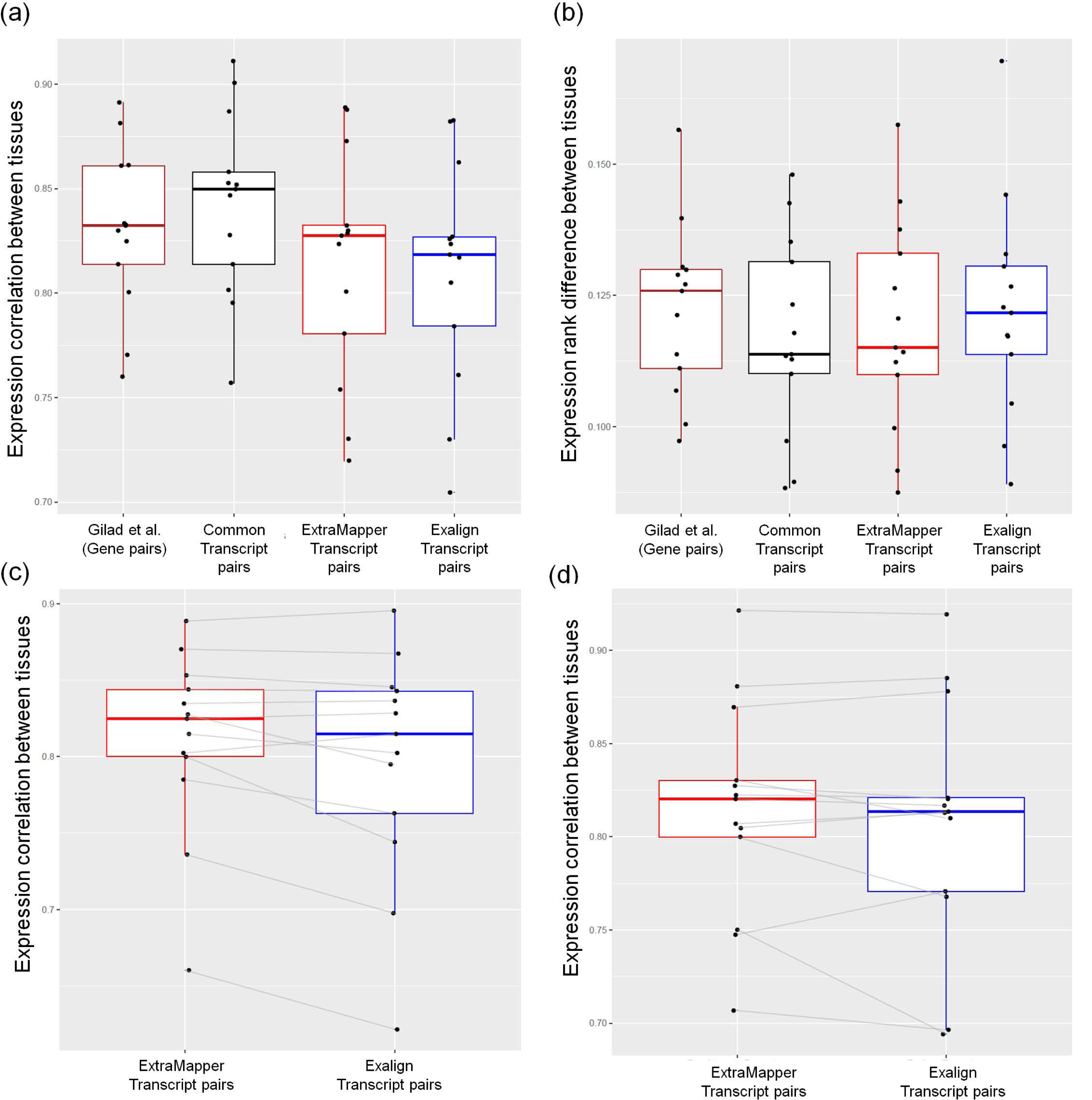
Reanalysis of tissue-matched human and mouse RNA-seq gene expression profiles. **(a)** Gene- and transcript-level expression correlation for tissue-matched human-mouse RNA-seq data. **(b)** Gene- and transcript-level expression rank difference for tissue-matched human-mouse RNA-seq data. **(c)** Gene- and transcript-level expression correlation for human transcripts where ExTraMapper and Exalign both report a single matching mouse transcript but their reported pairs are different from each other. **(d)** Similar plot to **(c)** but for mouse transcripts with a single matching human transcript. Each dot for each box plot represents a specific tissue type.

### A case study: Finding transcript-level mappings of the *BRAF-Braf* gene pair

Human *BRAF* gene is a proto-oncogene on chromosome 7 that encodes a RAF kinase (BRAF), which participates in the MAP kinase/ERK signalling pathway. Mutations in *BRAF* gene are shown to play important roles in multiple cancers including melanoma, long and colon cancers as well as in developmental disorders (37-39). *BRAF* is shown to be conserved among multiple organisms including fruit fly, worm and mouse. *Braf*, located on chromosome 6, is the mouse ortholog of human *BRAF* gene. According to Ensembl annotations (release 81), *BRAF* and *Braf* have 33 and 42 exons, respectively. The human ortholog spans 205 kb whereas the mouse one is around 122 kb long. Both genes have 5 transcript isoforms and 2 of them in each case are protein coding.

In Ensembl database, the orthology between *BRAF* and *Braf* genes is reported as a one-to-one relationship between two protein isoforms ENSP00000288602 and ENSMUSP00000002487 are encoded by the transcripts ENST00000288602 (BRAF-001) and ENSMUST00000002487 (Braf-001), respectively. **Fig. 6a** illustrates the best mapping of the exons of these two transcripts using pairwise coding exon similarities. Braf-001 has 4 more exons and codes for 38 more amino acids (aa) compared to BRAF-001 (22 vs 18 and 804 vs 766). Third exon of BRAF-001 with length 263 corresponds to a fusion of two exons in Braf-001 (lengths of 69 and 94) and the retention of the 100bp intron between them. Braf-001’s exon of length 35 is lost in human because of mutations in the donor and acceptor splice sites. Another Braf-001 exon with length 119 is also lost in human even though the splice sites are conserved. Also, BRAF-001 terminates one exon early compared to Braf-001. Overall, there are considerable differences between these two transcripts and their corresponding protein isoforms even though Ensembl orthology is defined through them. To the best of our knowledge, Ensembl gene-level orthology relationships are defined through pairing the longest protein isoforms of each organism even though they may not be the most similar ones.

**Figure 6.**
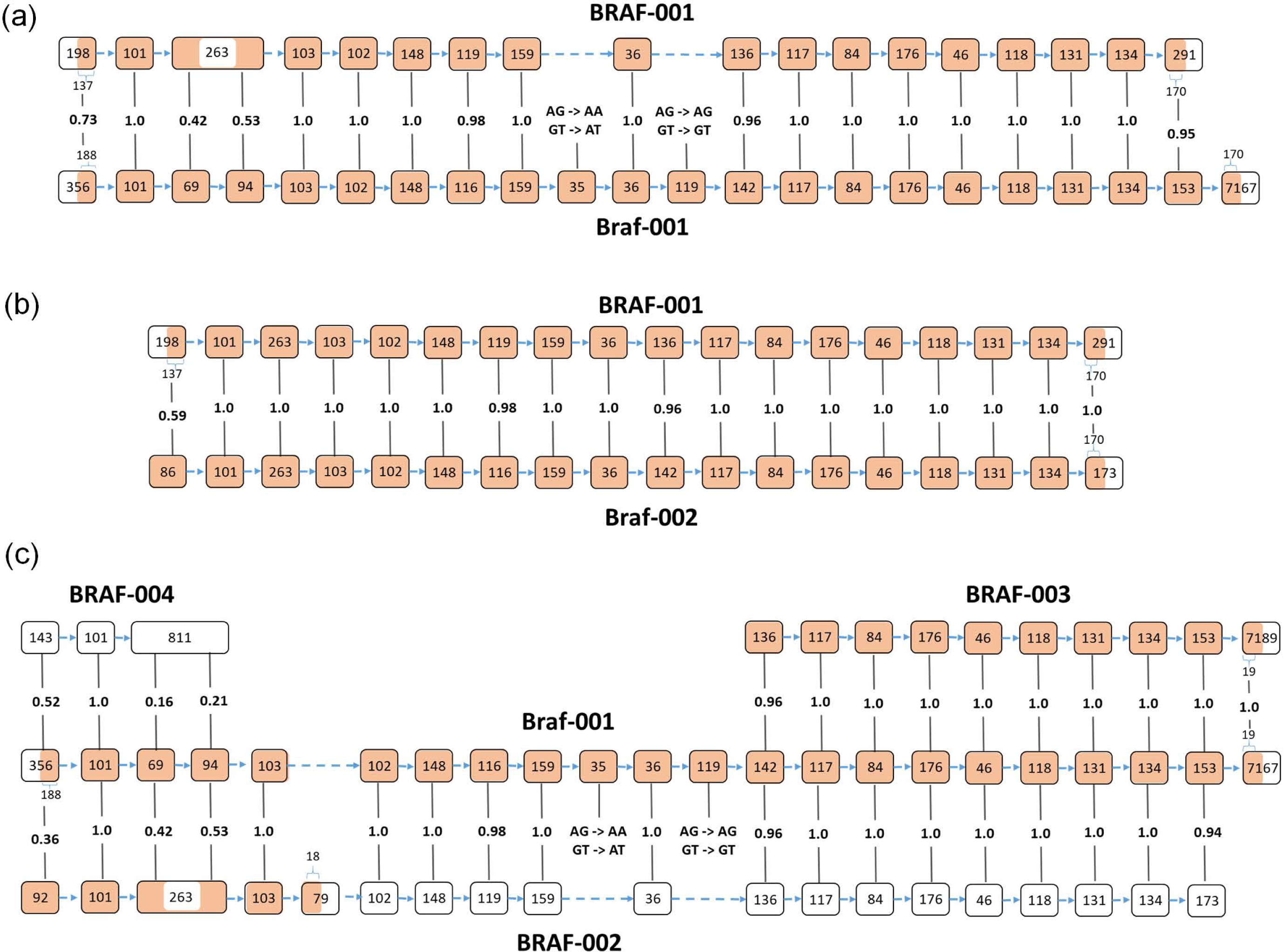
Transcript-level mappings for the *BRAF-Braf* gene pair. **(a)** Visualization of the transcript mapping between BRAF-001 and Braf-001 which is reported as the basis for orthology between the two genes by Ensembl. The coding similarity for this transcript pair is 0.86. **(b)** Visualization of the highest scoring transcript pair reported by ExTraMapper. The coding similarity for this transcript pair is 0.97. **(c)** Visualization of partial mappings from multiple human transcripts to the longest mouse transcript of Braf-001 with 22 coding exons including the part-coding first and last exons. Among these potential human transcripts that can be mapped to Braf-001, ExTraMapper assigns a coding similarity score of 0.19 and an overall similarity score of 0.82 to BRAF-002. For BRAF-003 which has a nearly perfect mapping to a part of Braf-001 both the coding and overall similarities are 0.62.

**Fig. 6b** illustrates the best transcript-level mapping identified with ExTraMapper, which links protein isoforms ENSP00000288602 and ENSMUSP00000099036 that are encoded by the transcripts ENST00000288602 (BRAF-001) and ENSMUST00000101497 (Braf-002), respectively. These two transcripts have the same number of exons and an almost perfect mapping between their exons except the first ones. ExTraMapper coding transcript similarity score for this pair is 0.97 out of 1 whereas it was 0.86 for the transcript pair reported by Ensembl gene orthology illustrating the need for systematic pairing at the transcript level. **Fig. 6c** demonstrates other suboptimal alignments found between the longest mouse transcript Braf-001 (protein coding) and three human transcripts BRAF-002 (nonsense mediated decay), BRAF-003 (protein coding) and BRAF-004 (retained intron). Notably, BRAF-002 contains a premature stop codon that limits the coding part to its first five exons, BRAF-003 is a near perfect match to the last 10 exons of Braf-001 initiated by an alternative first exon of length 136. BRAF-004 contains a large exon of length 811 which retained multiple intronic regions surrounding two exons in mouse. Overall, this case study provides us examples of complex exonic and splicing events that lead to the different transcripts with varying biotypes in human and mouse. Furthermore, it demonstrates the use of ExTraMapper in identifying such events as well as maximum similarity transcript pairings that are not readily available in any public dataset to the best of our knowledge.

## DISCUSSION

Gene regulation is one of the most fundamental processes that confer cellular identity and modulates the biological activities within a cell. However, most of the current analyses on gene regulation are carried out under the ’one gene -> one protein -> one function’ paradigm, thereby ignore the use of alternative promoters and alternative splicing in mammalian genes. Evolutionary studies that aim at identifying orthology relationships between organisms is not an exception to this trend and are also mostly focused on gene-level orthology. For example, HomoloGene (http://www.ncbi.nlm.nih.gov/homologene) is a system for automated detection of homologs among several completely sequenced eukaryotic genomes at the gene level. However, the 16k gene pairs that are orthologs between human and mouse span a diverse repertoire of over 200k total transcript isoforms more than 125k of which are protein coding. Current tools, including HomoloGene do not provide any information on which of these transcripts from mouse correspond to those in human in terms of their DNA sequence, protein sequence and function.

Here we described a novel method, ExTraMapper, which leverages sequence conservation between exons of a pair of organisms and identifies a fine-scale orthology mapping at the exon and then transcript level. ExTraMapper identified more than 250k exon, as well as 30k transcript mappings between human and mouse genomes. On several case studies, we demonstrated that ExTraMapper identifies exon fusions, splits and losses due to splice site mutations, and finds better transcript-level mappings compared to Ensembl orthology. Our results comparing human and mouse genomes showed that ExTraMapper finds mappings for more than 95% of exons that are involved in at least half of the annotated transcripts in either genome in agreement with previous studies that use a 20 times smaller subset of exons than used here. Compared to two existing exon mapping methods (OrthoExon and Inparanoid), ExTraMapper identified two to three times more perfect exon mappings (100% conservation) between human and mouse. By using both coding and overall conservation of an exon, we identified more than 30k mappings between part-coding exons and a large number of microexon mappings both of which were largely missed by the existing methods.

At the transcript level, ExTraMapper identified mainly one-to-one mappings using a greedy approach with multiple tie breakers. We reported 30,388 transcript mappings with a similarity score above 0.8 between 30,190 human and 30,150 mouse transcripts out of which 8,007 were perfect transcript mappings (100% conservation). We presented these transcript mappings for genes with known isoform-specific functions (*RARA, TP53, TP63*, and *TP73*) and created genome browser tracks for visualizing such mappings from either one of the compared genomes (UCSC visualizations for the *FOXP3-Foxp3* gene pair; **Supp. Fig. 10**). We also showed that a previous method that exclusively uses exon lengths in a Blast-like transcript alignment has a strong bias in mapping transcripts with larger number of exons and that ExTraMapper alleviates this bias resulting in mappings for transcripts from all length scales. For an orthogonal validation, we then used RNA-seq data from 13 different human and mouse tissues and compared the expression profiles of identified orthologous transcripts by Exalign and ExTraMapper. This analysis showed that transcript pairs identified by ExTraMapper have more correlated expression patterns compared to Exalign identified pairs. Further research directions include better annotation of transcript mappings found by ExTraMapper as well as its extension to include multiple mammalian genomes simultaneously.

We believe that ExTraMapper will have a great impact for translational sciences as it provides a dictionary for translating transcript-level information about gene expression and gene regulation from one organism to another. For example, one direction this translation can be done is from mouse models to human genome. This will be specifically useful for work with certain tissues or samples that are difficult to obtain from human donors or come in very limited quantities. Another important aspect of our work is that it allows us to identify which genes have highly conserved exon-intron structures and transcript repertoires between two organisms such as human and mouse. This information is important for understanding the extent of evolutionary differences with respect to specific gene functions and biological pathways. For instance, our GO term enrichment analyses showed that human genes that have all their protein coding transcripts map perfectly to a corresponding mouse transcript are enriched in certain developmental processes such pattern specification process and regionalization. On the other hand, genes that have none of their coding transcripts map to any mouse transcript are enriched in response related processes such as defense response to other organism suggesting higher level of divergence between human and mouse in their immune system related transcripts.

## MATERIAL AND METHODS

### Publicly available datasets and methods

#### Gene annotations and orthologous pairs

We download and use gene annotations and orthology information from Ensembl database release 81(40). For human genome, we use the GTF file Homo_sapiens.GRCh38.81.gtf.gz and the corresponding reference genome build of hg38. For mouse genome, we use the GTF file Mus_musculus.GRCm38.81.gtf.gz and the corresponding reference genome build mm10. For the orthology information of gene pairs between human and mouse genomes, we use the files ensembl_mart_81/hsapiens_gene_ensembl__homolog_mmus dm.txt.gz and ensembl_mart_81/ mmusculus_gene_ensembl homolog_hsap dm.txt.gz. These files provide orthology pairings for 16,711 genes, 15,846 of which are annotated as protein coding genes in both organisms. Almost all the remaining orthologous gene pairs code for various types of RNA and are between single-exon, single-transcript genes which are trivial cases for our interest. Summary of exon and transcript-level annotations for the 15,846 orthologous gene pairs are presented in **Table 1**.

#### Conservation scores between human and mouse

We use Multiz alignments provided as MAF (Multiple Alignment Format) files and PhastCons to compute a conservation score for each human and mouse exon (41-43). We filter the MAF files downloaded from http://hgdownload.cse.ucsc.edu/goldenPath/hg38/multiz7way/hg38.7way.maf.gz for human genome and from http://hgdownload.cse.ucsc.edu/goldenPath/mm10/multiz60way/maf/ for mouse genome by using *maf_parse* to only keep human to mouse and mouse to human alignments. We then use *phastCons* twice, once to obtain tree models and once to generate wig files with conservation scores at single base pair resolution. Using these scores, for each exon in either organism, we compute average conservation scores (i) for the whole exon body, (ii) for the coding portion of the exon and, (iii) for the acceptor and donor sites of the exon.

#### Exon mappings from OrthoExon (20)

We download mappings between united exons described in Fu et al from http://tdl.ibms.sinica.edu.tw/OrthoExon/download.html. Since the exon coordinates in these mappings belong to earlier reference genome versions (hg18 for human and mm9 for mouse), we first use *liftOver* to convert these coordinates and then use *bedtools intersect* to find corresponding exons in hg38 and mm10 genome builds. **Fig. 3** provides a summary of the resulting exon mappings.

#### Exon mappings from protein isoforms using Inparanoid (22)

We download mappings between orthologous exons described in Zhang et al from http://genomebiology.com/content/supplementary/gb-2009-10-11-r120-s5.zip. Since the exon coordinates in the mappings provided in Dataset_s2_Human_and_mouse_orthologous_exons_list.tsv belong to earlier reference genome versions (hg18 for human and mm8 for mouse), we first use *liftOver* to convert these coordinates and then use *bedtools intersect* to find corresponding exons in hg38 and mm10 genome builds. **Fig. 3** provides a summary of the resulting exon mappings.

#### Transcript mappings from Exalign (16,17)

We download Exalign software from the link http://159.149.160.51/exalign/Download/exalign.tar.gz. We then parse the transcript information for only protein coding transcripts from Ensembl database (release 81) for human and mouse in the input format desired by Exalign. We use “*exalign -O <organism>*.*rf –freq*” to compute exon length frequency files required by Exalign for each organism using all their genes. Then, for each orthologous gene pair, we create a query/database file pair using only the transcripts for that gene pair. We then run *exalign* with default parameters to find transcript mappings from human to mouse and from mouse to human using exon length frequencies generated from all genes.

#### Tissue specific RNA-seq data, and expression profile analysis

We download the human and mouse tissue specific RNA-seq data (35) from ENCODE project site in FASTQ format (23). For our analysis we first map the human and mouse tissue specific RNA-seq reads on GRCh38 and GRCm38 genome assembly, respectively, using Hisat2 (44). We then use *featureCounts* (45) to generate the initial read counts for gene, transcript and exon-wise Ensembl genomic annotations (24) for each tissue per organism from the mapped reads. We next follow a similar protocol as described by Gilad et al. (36) to account for the sequencing study design batch effect using “ComBat” function from “sva” package to get the final gene, transcript and exon-wise expression values for downstream analysis.

### Detailed description of ExTraMapper

#### Computation of pairwise exon similarities

From the 15,846 orthologous gene pairs, we first extract all exons from all annotated transcripts for both human and mouse genomes. We use *liftOver* chains between the two organisms to find, for each exon coordinate in one organism, the corresponding genomic coordinates in the other organism. To allow for non-perfect exon mappings, we use multiple settings for the “-minMatch” parameter (0.9, 0.95 and 1.0) of *liftOver* which sets the minimum ratio of bases that must remap to the target organism (default value is 0.95). Also, to capture potential one-to-many mappings, we use the “-multiple” option of *liftOver* which allows outputting multiple target regions per input region. For each gene pair ***g***, 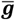 for each exon ***e***_***i***_ of the gene ***g*** in one organism, we find the exons of the gene 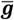 rom the other organism that intersect with the lifted over coordinates of ***e***_***i***_. For each possible pairing of the ***e***_***i***_ from ***g*** and 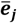 from 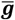 we compute a similarity score using:

i. the amount of overlap between the lifted over ***e***_***i***_ coordinates and the original 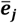 coordinates,
ii. the ratio between the original length and the lifted over length of ***e***_***i***_.

We compute the same score for the 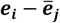 pair by reversing the order of the two organisms and take the maximum score between the two orderings. More formally, let ***lf***(***e***) denote the lifted over coordinates and |***e***| denote the length of the exon ***e***. Then, we define the similarity for the exon pair ***e***_***i***_, 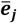 as:

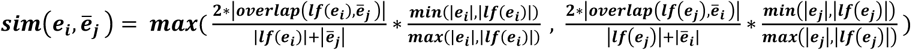

This score is zero for any two exons that do not overlap after both ways of liftOver and it is proportional to the percentage of overlap for overlapping exon pairs. The second terms on each side of the *max* operator account for cases in which the lifted over coordinates correspond to shorter or longer regions with respect to the original exon coordinates and is 1 for length preserving liftOver operations which are usually the case. A similarity score of 1 corresponds to perfect conservation of length and overlap for an exon between human and mouse genomes within an orthologous gene pair.

We calculate the above similarity over the whole exon body regardless of whether the exon is fully coding, partially coding or non-coding. To compute a coding similarity counterpart of this overall exon similarity, we analyze three cases of exon coding type separately. For the case of fully coding exons, the overall similarity equals coding similarity. For non-coding exons the coding similarity is trivially undefined and is set to zero. For the case of partially coding exons, we compute coding similarity score using only the coding portion of an exon. We use the coding portions of each partially coding exon for the liftOver between the organisms and then for the length and overlap calculations. We then use these values in the same formulation as above to compute the coding similarity score. We store both the overall and coding similarities, and use them in overall and coding similarity calculations for transcript pairs and for exon-level mappings.

#### Computation of pairwise transcript similarities

For each orthologous gene pair ***g***, 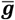 for each pair of transcripts ***t***_***i***_, 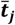 such that ***t***_***i***_ ∈ ***g***, and 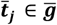 we compute two similarity scores (i.e., overall and coding) using the similarities of the conserved exons between the two organisms. Computation of these similarity scores in non-trivial since an exon of one organism sometimes overlaps with several other exons of the other. Accordingly, there are multiple different ways to compute the transcript level similarities. This problem bears resemblance to sequence alignment but exons are the units aligned instead of nucleotides. Therefore, one solution is to use ***dynamic programming*** to find the exon alignment with possible gaps using strictly similarity scores (zero or greater) and a gap penalty of zero. Another option is to use all exon pairs that have an exon similarity score of at least a given similarity threshold for the transcript-level similarity calculation. The third way to compute the transcript similarity is to ***greedily*** select the “most” similar exon pair between the two transcripts, match the two exons to each other, remove them from the set of exons to be paired and repeat this process until there is no exon pair with a similarity score of at least a given threshold. For this greedy approach, determining the “most” similar exon pair at each step requires tie break conditions as ties are quite common. For this purpose, we first look at the coding similarity of each exon pair and if there are multiple exon pairs with the same similarity score we then break the tie by the overall exon similarity. If there are still ties, we pick one exon pair and report it and then repeat the process. Note that this will not exclude unpicked exon pairs which will be reconsidered in the next iteration unless one element of the exon pair is already reported.

Once a set of one-to-one and non-contradicting exons mappings ***E***_***m***_are found for a transcript pair ***t***_***i***_, 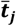, we can then define the similarity score as:

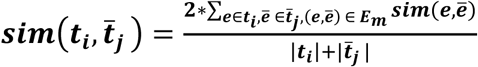

Note that for partially coding exons, we take ***sim(e***, 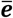) s the maximum of overall exon similarity and coding exon similarity. We use all exons, including non-coding exons, for the overall pairwise transcript similarity. For the coding transcript similarity, we use only the exons that are either fully coding or partially coding for both exon similarity and transcript length calculations in the above equation. These transcript similarity scores range between 0 and 1, with a similarity score of 1 indicating perfect conservation of a transcript between human and mouse genomes including the transcript length and exon identity.

#### Extraction of transcript-level mappings between human and mouse

Once we compute pairwise transcript similarities for each gene pair, we get a transcript similarity matrix ***T*** that is *n* × *m* where *n* and *m* correspond to number of transcripts of the two genes in the orthologous gene pair. Each transcript from one organism potentially has similarity to multiple transcripts from the other organism leading to one-to-many or many-to-many relationships among the transcripts of the two organisms. We use a greedy method to extract a set of transcript mappings which mostly includes one-to-one mappings that do not conflict with each other. Note that, it is possible to extract such set of mappings with highest total similarity score by solving the maximum weight bipartite matching (MWBM) which can be done in polynomial time. However, MWBM prefers multiple suboptimal mappings (e.g., two mappings with score of 0.6 and 0.5) to a perfect mapping of score 1 which conflicts with the two suboptimal mappings. This feature, even though desirable in some scenarios, is undesirable for our purpose which is to find a set of transcript mappings, even though it may be a small set, that are highly conserved between the two organisms. Therefore, we greedily select the transcript pair with the highest similarity score first, accept this transcript mapping into our results set and then set the corresponding row and column in matrix ***T*** to zero to ensure no conflicting mappings will selected in the later iterations. We repeat this process until no more non-zero entries are left in ***T***. We also post-process the set resulting set of mappings and discard those transcript pairs with a similarity score below a given threshold (at least 0.8) to ensure that only highly conserved pairs are reported.

Since the above described method of extracting transcript mappings can include ties, such as two pairs with the same exact score in ***T***, in the process, we break these ties in the best way possible to favor pairs that we believe are more similar. We list the criteria for breaking these ties and their ordering in **Fig. 1**. Briefly, we first look at the coding similarity between transcripts to pick a single transcript pair. In case of a tie for this criterion, we resort to overall transcript similarity to break the tie among the tied pairs from the previous step. If there is another tie, we pick among the tied pairs the transcript pair that has the smallest difference between the numbers of coding exons of the two transcripts. If there are still tied pairs after all the previous criteria, we try to break it by the difference between the numbers of overall exons of the two transcripts. Lastly, we pick and report all transcript pairs that are tied in all the four criteria we checked because all these pairs have equivalent value for our purposes.

## ACKNOWLEDGEMENTS

We would like to thank Wen-chang Lin for making their OrthoExon database available for batch download.

## SUPPLEMENTARY DATA FILES

**Supplementary Data File 1 contains Supplementary Figures 1-10 (pdf). Supplementary Data File 2 contains Table 1 (xlsx)**.

**Table 1**. Summary of exon and transcript annotations from Ensembl (release 81) for human and mouse genomes.

**Supplementary Data File 3 contains Table 2 (xlsx)**.

**Table 2**. Transcript-level mappings reported by ExTraMapper for three human mouse gene pairs *TP53-Trp53, TP63-Trp63* and *TP73-Trp73*.

**Supplementary Data File 4 contains Table 3 (xlsx)**.

**Table 3**. Accession IDs for human and mouse RNA-seq data.

