## Supplementary material for "ExTraMapper: Exon- and Transcript-level mappings for orthologous gene pairs": Supp Info

**Supplementary Information**

### SUPPLEMENTARY FIGURES

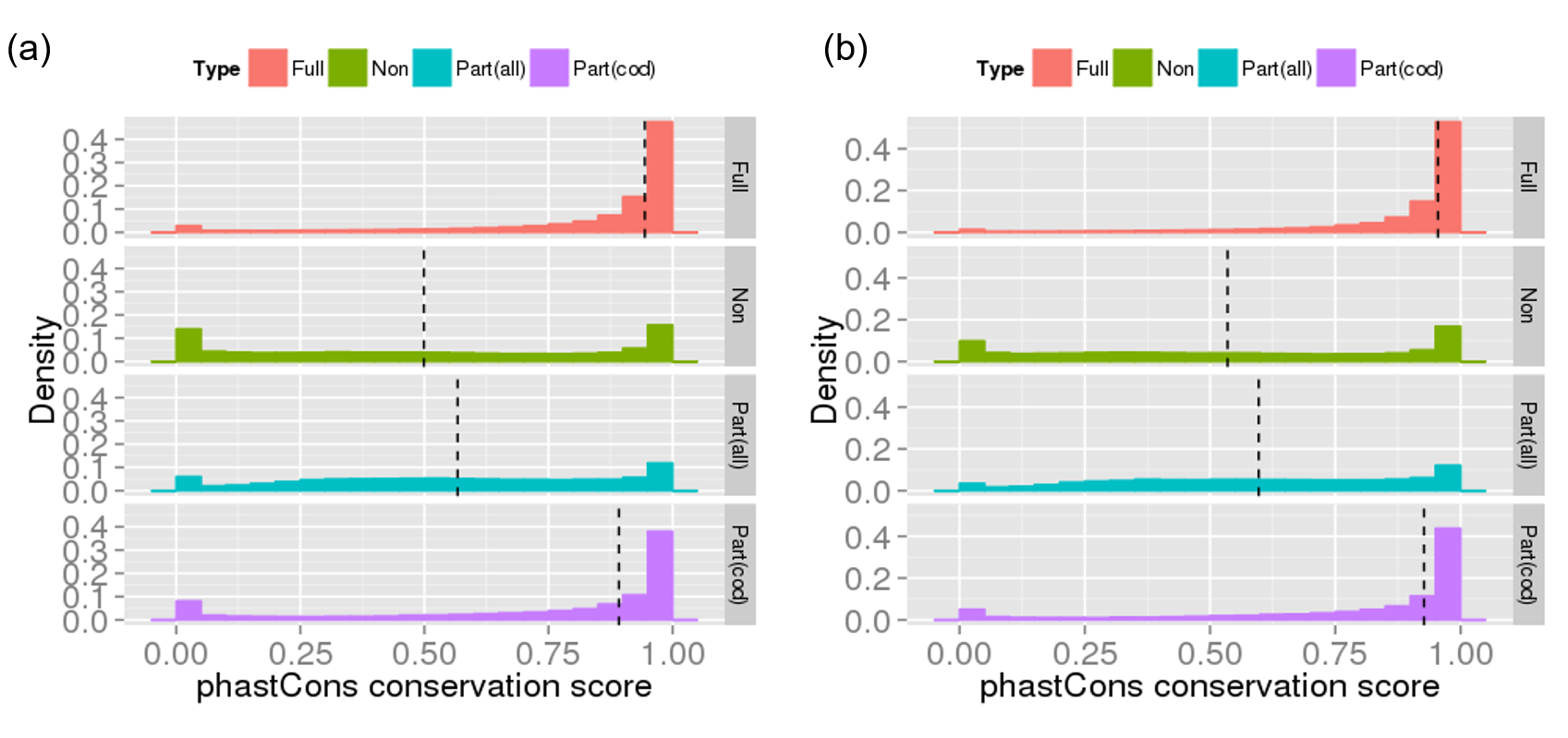

Supplementary Figure 1:

Distribution of evolutionary conservation scores for different coding types for (a) human and (b) mouse exons. The conservation scores are computed using PhastCons (Methods). From top to down the different types in each figure are: fully coding exons, non-coding exons, partially coding exons and only coding parts of partially coding exons.

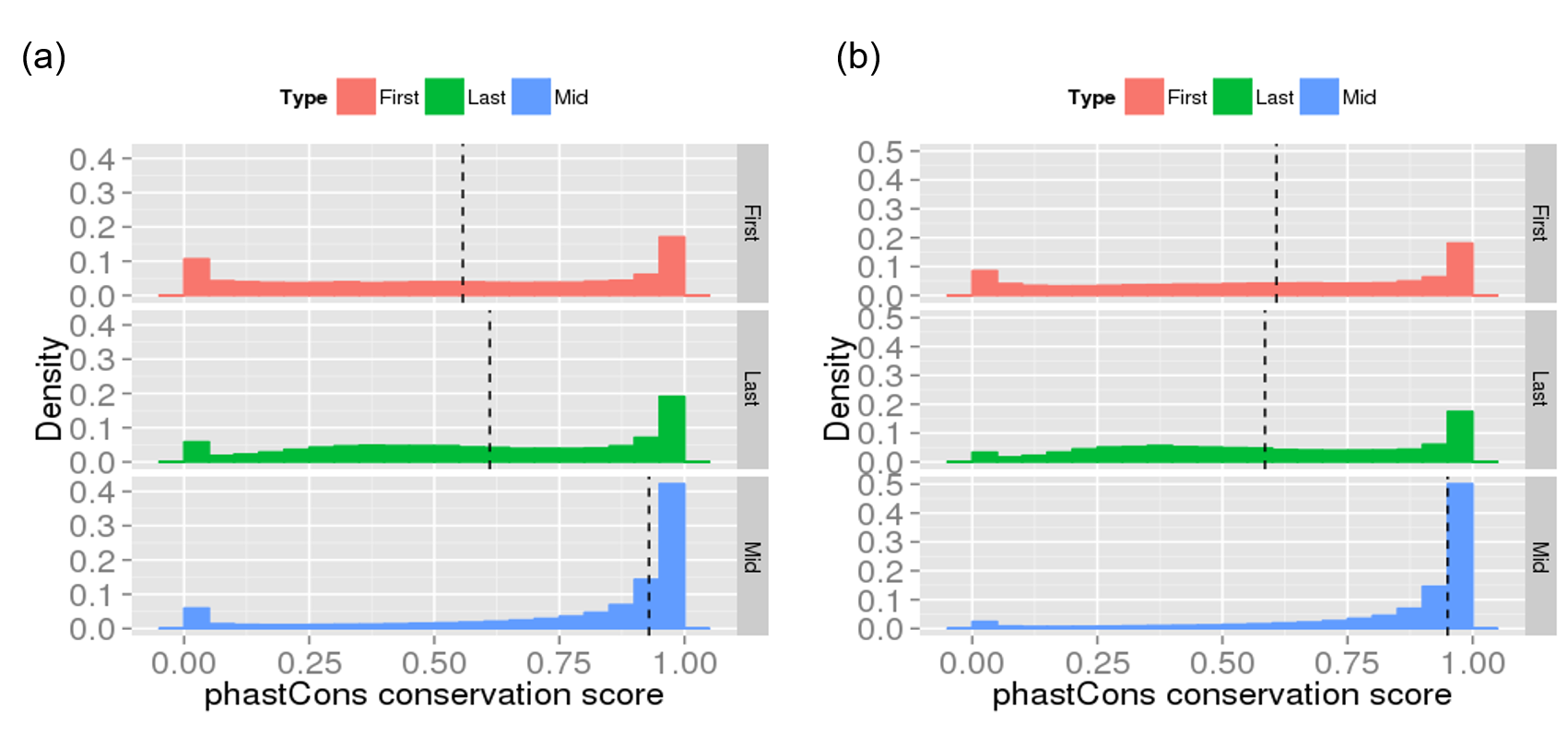

Supplementary Figure 2:

Distribution of evolutionary conservation scores for different exon position types for (a) human and (b) mouse exons. The conservation scores are computed using PhastCons (Methods). From top to down the different types in each figure are exons that are always: first, last and middle of each transcript.

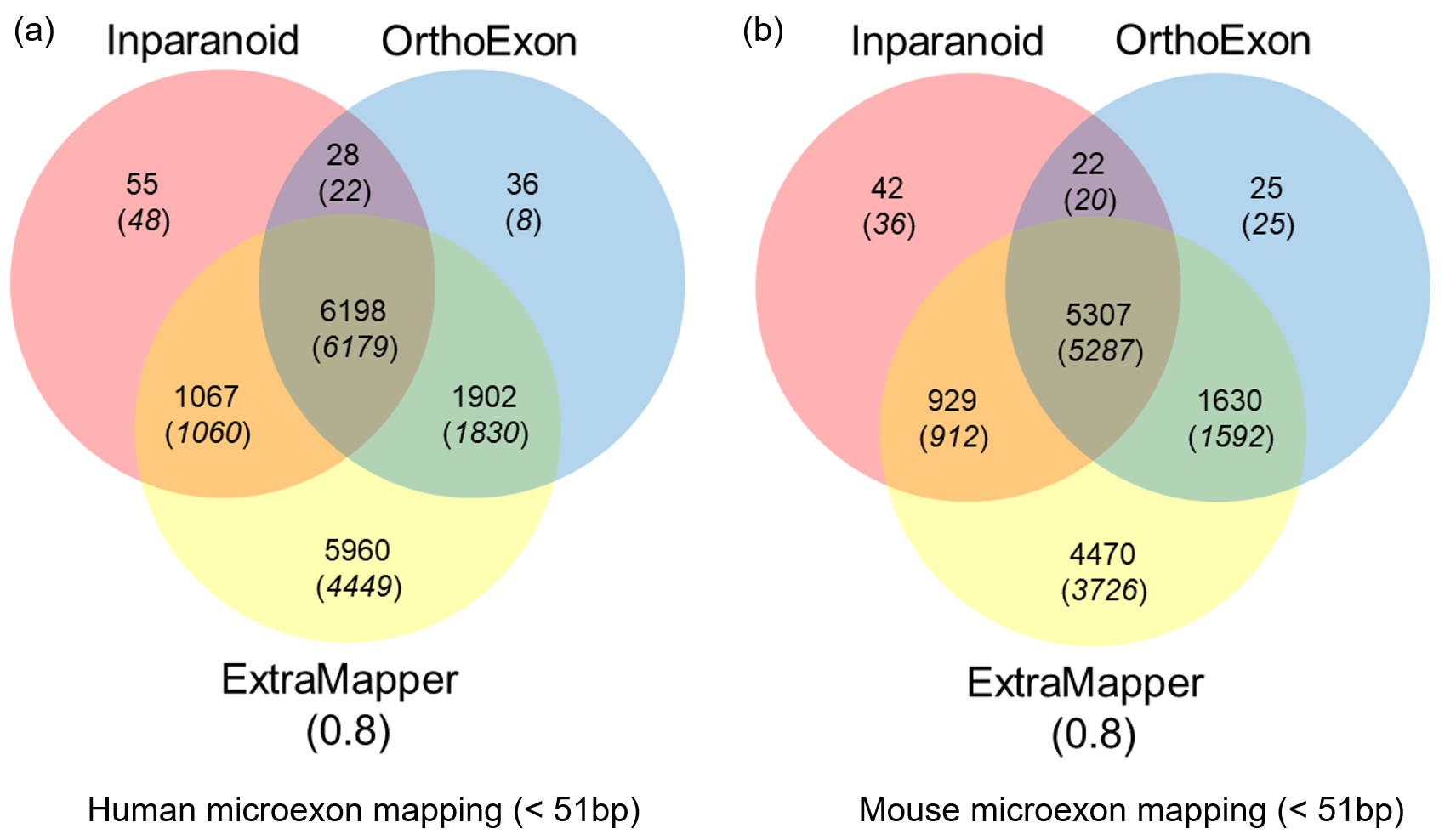

Supplementary Figure 3:

Venn diagrams showing intersections among human-mouse exon mappings reported by three different methods (ExTraMapper, OrthoExon and Inparanoid) for short exons (or microexons if defined broadly) that are less than 51-bp long. We use an exon similarity score threshold of 0.8 for ExTraMapper to determine the set of exon mappings. Results for the number of (a) human and (b) mouse microexons are shown separately.

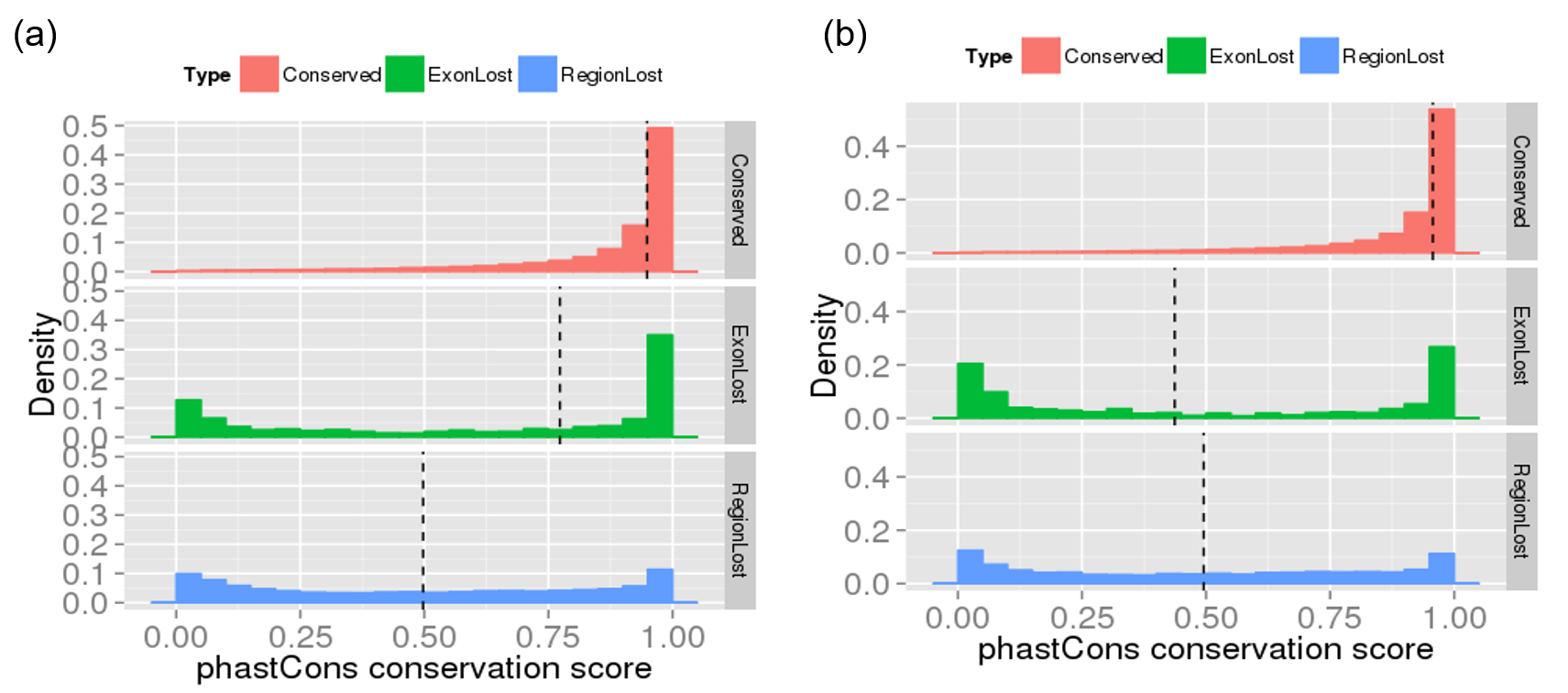

Supplementary Figure 4:

Distribution of evolutionary conservation scores for different exon classes determined by ExTraMapper for (a) human and (b) mouse exons. The conservation scores are computed using *PhastCons*. From top to down the different types in each figure are exons that are classified as: conserved between human and mouse, region conserved but lost exon function in the other organism and exon regions that are lost in the other organism.

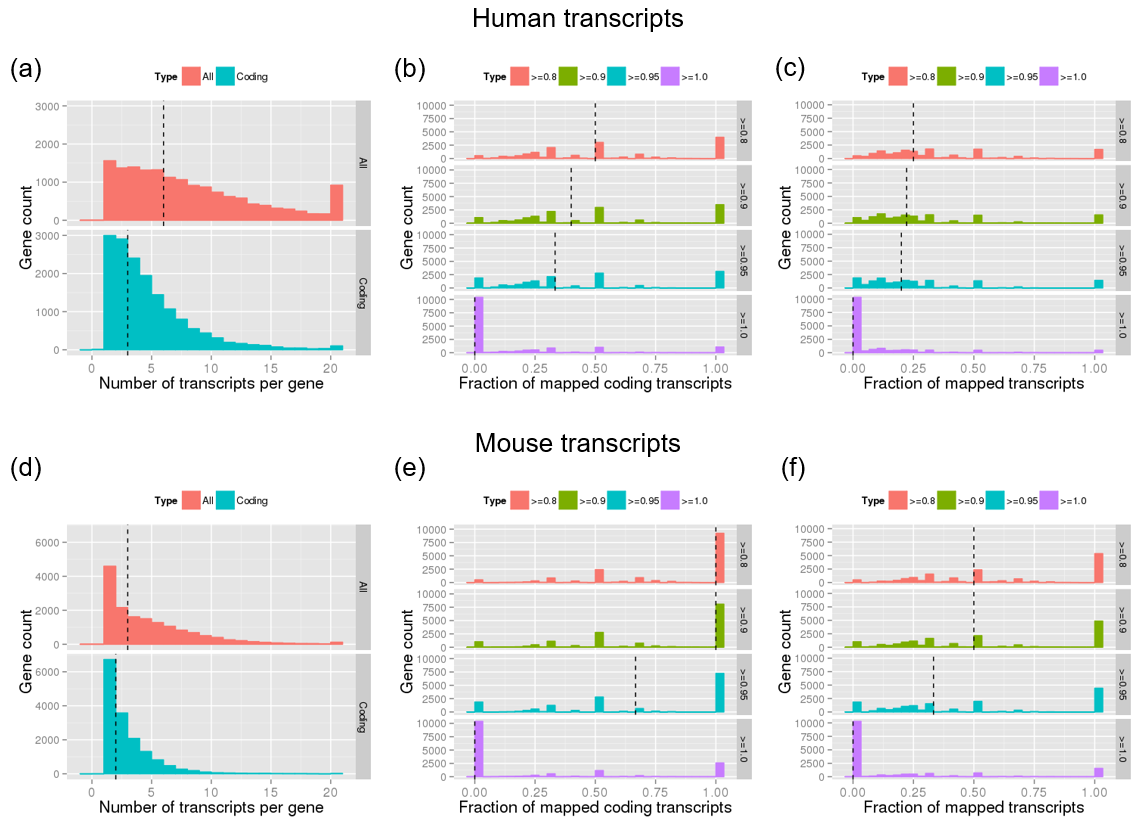

Supplementary Figure 5:

Summary of transcript-level statistics and ExTraMapper mapping results for human (top row) and mouse (bottom row). **(a,d)** The histograms of the number of transcripts per gene. **(b,e)** The histograms of the fraction of coding transcripts that are mapped when different stringency cutoffs for mapping similarity are used. A fraction of 1 indicates all transcripts of a gene are mapped and 0 indicates none are mapped. **(c,f)** Similar histograms using all transcripts. Dashed vertical lines correspond to median of the distribution for each plot.

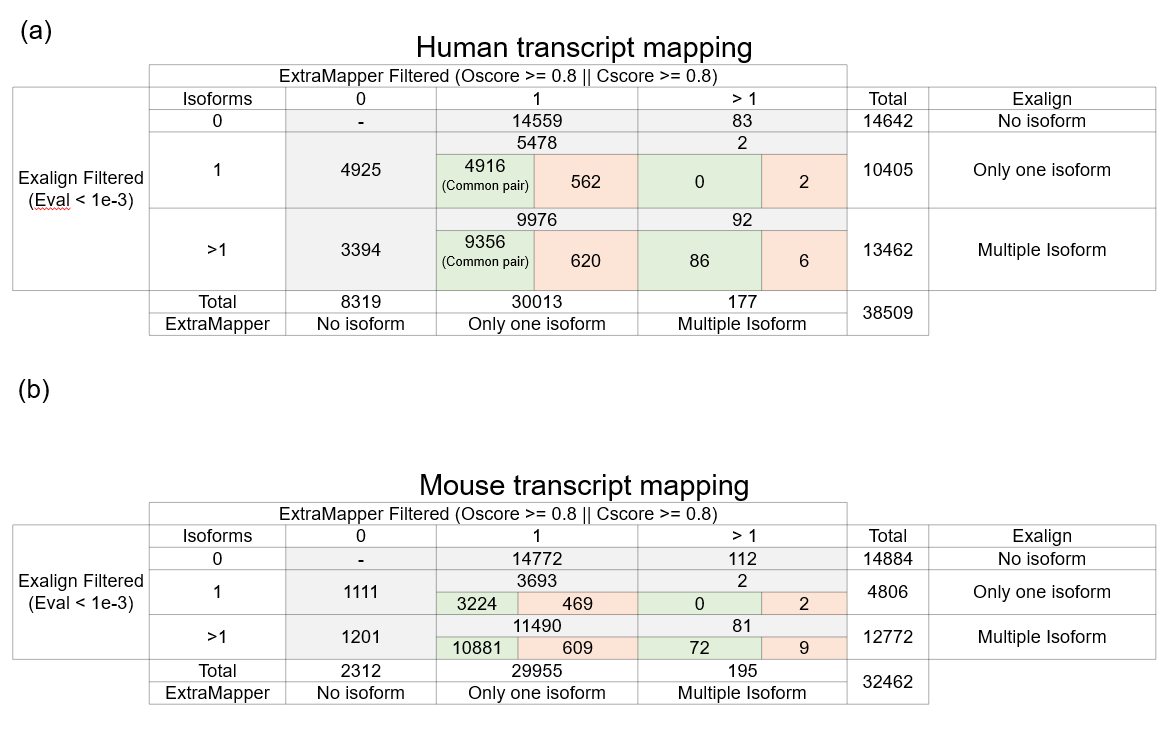

Supplementary Figure 6:

Comparison of transcript mapping in **(a)** human and **(b)** mouse by ExTraMapper and Exalign programs. The total number of human transcripts that are exactly mapped to only one mouse transcript is 30,013 for ExTraMapper (score >0.8) while this number was only 10,405 for Exalign. ExTraMapper only reports 177 human transcripts that map to more than one mouse isoform, whereas Exalign reports 13,462 such human transcript. A similar trend can also be seen when numbers are computed with respect to mouse transcripts.

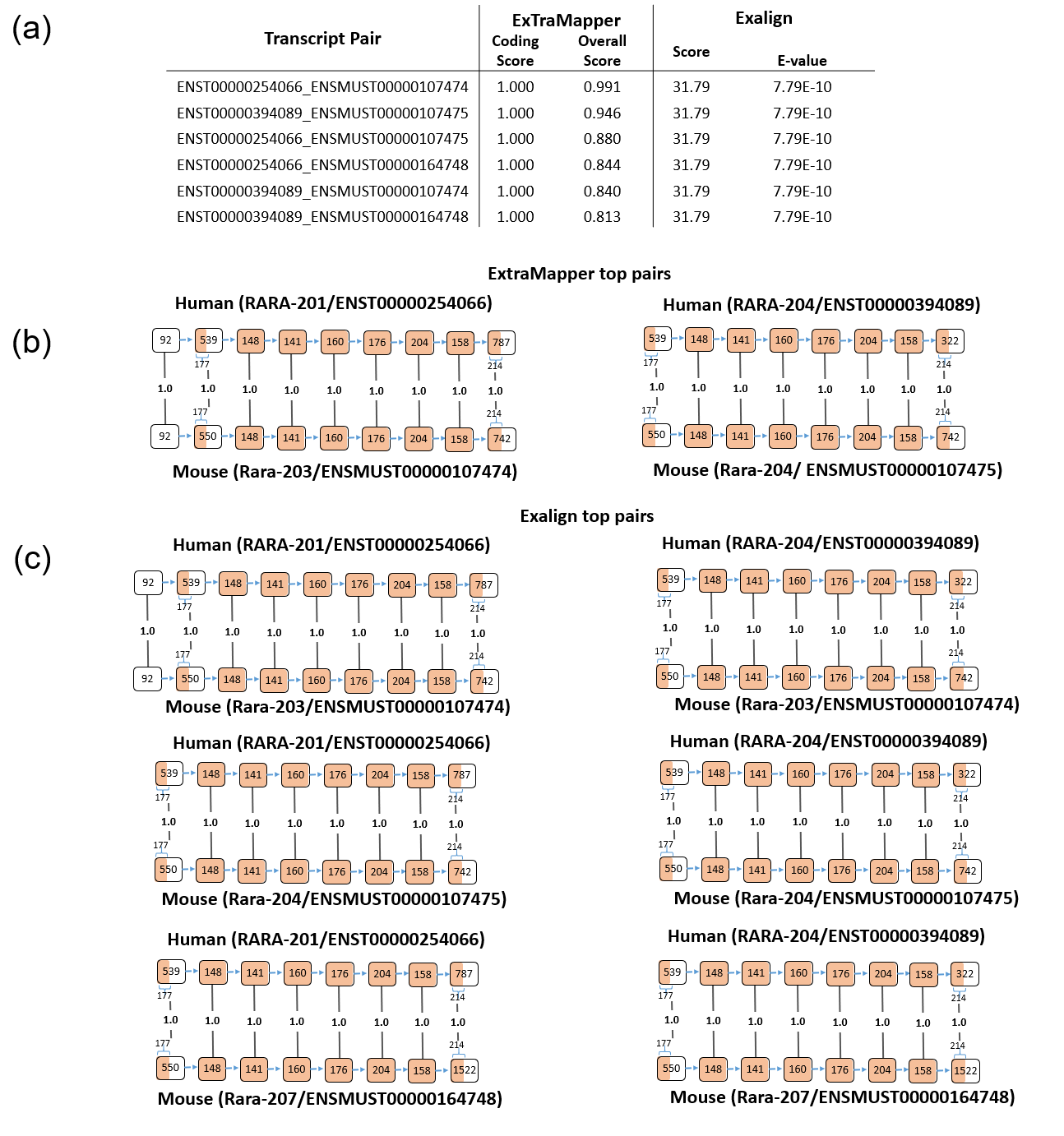

Supplementary Figure 7:

**(a)** Retinoic acid receptor alpha (*RARa*) transcript isoform mapping using ExTraMapper and Exalign programs between human and mouse. **(b)** Out of six possible pairs among two human and three mouse isoforms for this gene, ExTraMapper breaks ties using the similarity between non-coding portions of these transcripts to report two one-to-one transcript mappings. **(c)** Exalign fails to break the ties and reports the same score for every combination.

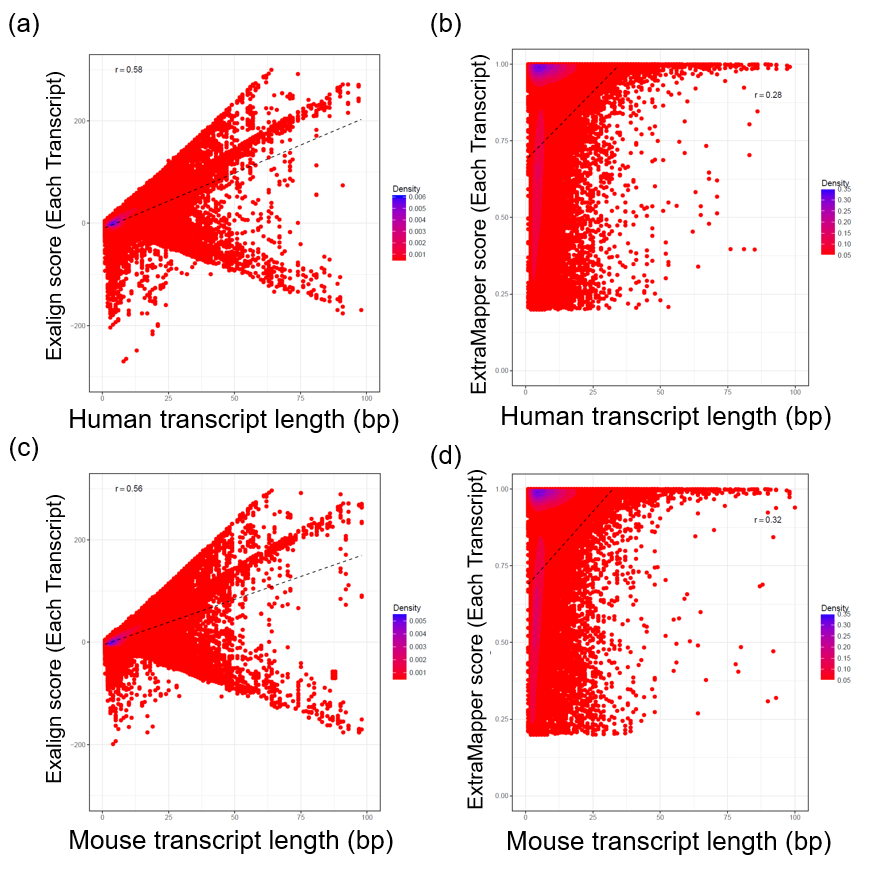

Supplementary Figure 8:

The density plots of the Exalign alignment and the ExTraMapper similarity scores for all the transcript pair for each gene are plotted for **(a)** human transcripts using Exalign **(b)** human transcripts using ExTraMapper **(c)** mouse transcripts using Exalign **(d)** mouse transcripts using ExTraMapper. Transcript length indicates the number of coding (either fully or part-coding) exons of a transcript. Pearson correlation is reported for each figure together with a linear fit.

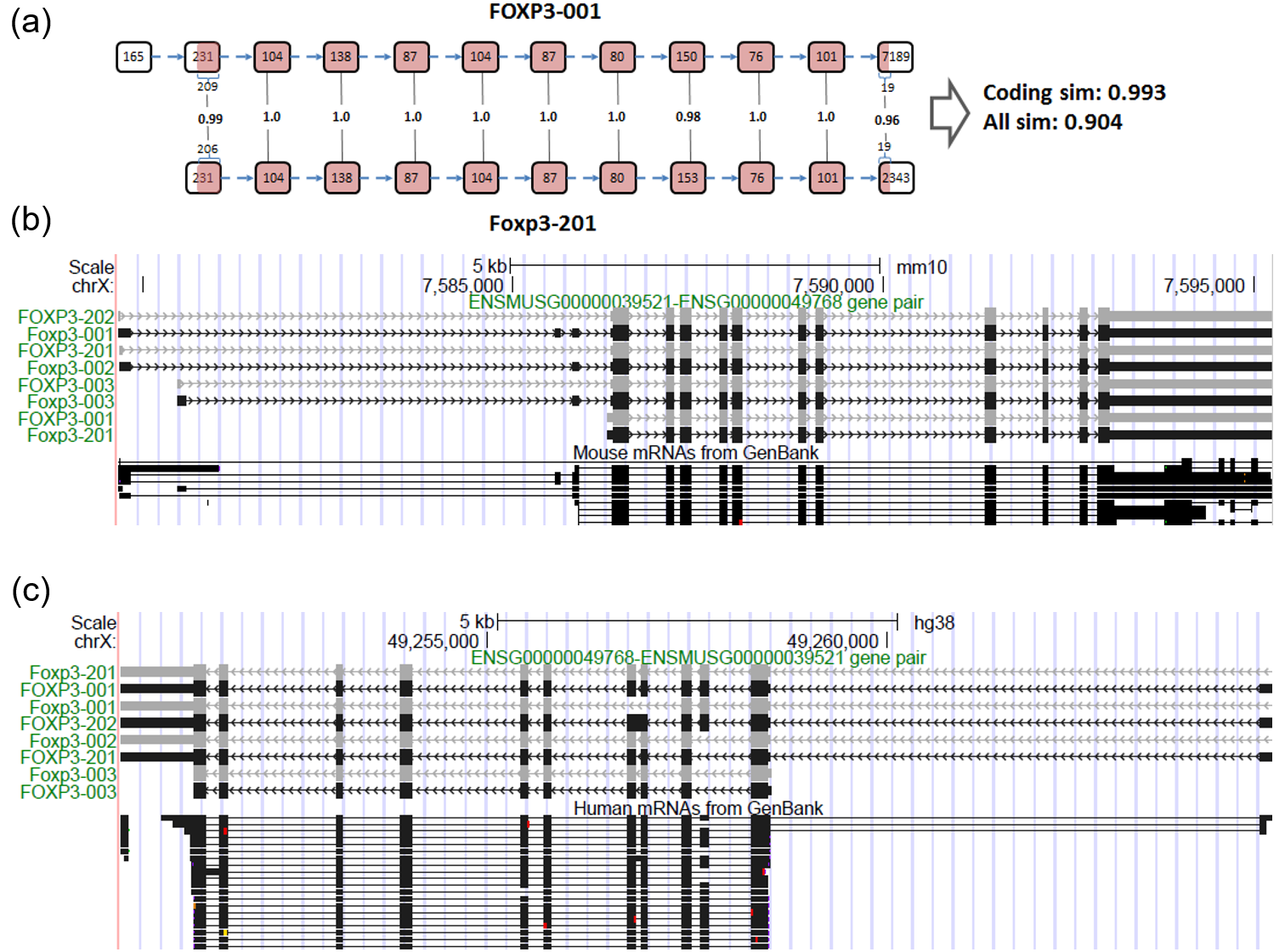

Supplementary Figure 9:

Transcript mappings for *FOXP3-Foxp3* gene pair. (a) The most similar transcript pair for this gene pair visualized using squares with exon lengths in base pairs and filled squares for protein coding regions. Arrows indicate the direction of or transcription and vertical connectors report the coding similarity between the vertically aligned exons one from human (top) and one from mouse (bottom). (b) UCSC mouse genome browser (mm10) snapshot of four pairs of transcripts that are mapped by ExTraMapper. Each human transcript (*FOXP3*) is followed by the corresponding mouse transcript (*Foxp3)*. All exons are shown for the mouse transcript whereas for human transcript only the exons that are mapped to a mouse exon are drawn. (c) UCSC human genome browser (hg38) snapshot which shows the same transcript mappings but from the view of all human exons and only the mouse exons mapped to those human exons.
